# Spatio-temporal brain invasion pattern of *Streptococcus pneumoniae* and dynamic changes of the cellular environment in meningitis pathogenesis

**DOI:** 10.1101/2023.10.30.564671

**Authors:** Kristine Farmen, Miguel Tofiño-Vian, Katrin Wellfelt, Lars Olson, Federico Iovino

## Abstract

*Streptococcus pneumoniae* (the pneumococcus) is the major cause of bacterial meningitis globally, and pneumococcal meningitis is associated with increased risk of long-term neurological sequelae. These include several sensorimotor functions that are controlled by specific brain regions which, during bacterial meningitis, are damaged by the vast neuroinflammation and bacterial toxins. Little is known about the invasion pattern of the pneumococcus into the brain. Using a bacteremia-derived meningitis mouse model, we combined 3D whole brain imaging with brain microdissection to show that all brain regions were equally affected during disease progression, with pneumococci in close association to the microvasculature. In the hippocampus, the invasion provoked a dynamic microglial response, while the dentate gyrus showed a significant loss of neuroblasts. Our results indicate that, even before symptom occur, the bacterial load throughout the brain causes neuroinflammation and cell death, a pathological scenario which ultimately leads to a failing regeneration of new neurons.

## Introduction

Pneumococcal meningitis, the inflammation of the meninges caused by the Gram-positive bacterium *Streptococcus pneumoniae* (the pneumococcus), is a medical emergency, with a mortality ranging between 18-30%, depending on the geographical region, with half of survivors suffering from long-term neurological sequelae, most commonly hearing loss, motor disabilities and cognitive impairments [1]. In the majority of pneumococcal meningitis cases, invasion into the brain and meninges follows a hematogenous route of infection, although translocation from the nasopharynx through the cribriform plate or through direct implementation in the brain following trauma or surgery does occur [2–4]. The presence of *S. pneumoniae* in the brain causes a vast inflammatory response, with release of reactive oxidative species and matrix metalloproteinases that augments pathological progression [5]. A consequence of the neuroinflammation is the increased intracranial pressure due to edema, which is linked with increased risk of cerebral infarction and ischemic injury in patients [6]. This, together with the direct toxicity of pneumococcal virulence factors, cause neuronal damage and death [2, 7].

Experimental models of pneumococcal meningitis have shown that S*. pneumoniae* penetrates the blood-brain barrier (BBB) in a transcellular pathway involving the platelet endothelial adhesion factor (PAF) receptor, platelet endothelial adhesion molecule (PECAM)-1 and the polymeric immunoglobulin receptor (pIgR) on brain vascular endothelial cells that binds the bacterial phosphorylcholine, the pneumococcal protein PspC and pilus-1, respectively [8, 9]. A paracellular pathway involving PECAM-1 has been observed, as well as BBB breakdown in patients, which further facilitate paracellular route of entry [10]. The pneumococcus has been shown to initially invade through the BBB of vessels close to the subarachnoid space in early stages of disease and, in later stages, to reach the inner brain through the choroid plexus, where it can then likely enter the cerebrospinal fluid [11]. This is in contrast to *Streptococcus suis,* where the choroid plexus has been reported as the main point of entry, while histopathological examination from *Neisseria meningitidis* meningitis show equal bacterial entry through both the microvasculature and the choroid plexus [12]. Even though the molecular interactions between pneumococci and brain endothelial cells have been characterized, it is still poorly understood whether pneumococcal entry into the brain occurs preferentially in certain brain regions [2, 10]. When it comes to cellular invasion, *S. pneumoniae* was recently shown to bind β-actin exposed on the neuronal plasma membrane through its pilus-1 tip protein RrgA, and through this interaction pneumococci invade neurons, ultimately causing neuronal death through cytoskeleton disruption [13]. Pneumococci produce H_2_O_2_, and together with the pore forming toxin pneumolysin contribute to the cellular death of neurons and astrocytes, while at the same time inhibiting the motility of microglia, thus reducing phagocytosis [14]. Microglia sense *S. pneumoniae* through extracellular receptors such as Toll-like receptors 2,3,4, and 9, as well as through intracellular receptors including nucleotide-binding oligomerization domain-like receptors [15, 16]. This recognition causes the release of extracellular cytokines and chemokines, which drives the subsequent infiltration of neutrophils into the brain [17]. While innate immune cells are capable of phagocytosing the bacterium, this clearance is ineffective, causing an upheld proinflammatory state, release of oxidative species, and increased neuronal damage [18]. Interestingly, neuroinflammation and neuronal damage can also occur without bacterial invasion into the brain. In fact, Orihuela et al, showed that, during bacteremia with no pneumococci present in the brain parenchyma, microglia were activated and neuronal damage in the hippocampal dentate gyrus was detected [19]. In patients that died of bacterial meningitis, apoptotic neurons in the dentate gyrus of the hippocampus were observed in the majority of cases [20]. The subgranular zone (SGZ) of the dentate gyrus represents one of the two main neurogenic niches in the adult brain. Newly generated cells in the SGZ can become integrated in the neuronal circuits involved in cognition, memory, and learning. Damage to these pathways has been shown to cause behavioral impairment [21, 22]. This has also been observed in experimental pneumococcal meningitis, where caspase-3 dependent apoptosis in the hippocampus was linked with learning deficiencies in the rodents [20]. Thus, the observed neurological sequelae in patients are a consequence of several pathophysiological processes and dependent on the severity and the brain regions affected.

While treatment with anti-inflammatory drugs in rodents attenuates brain damage, and dexamethasone treatment in patients has shown to be beneficial in terms of outcome [23, 24], there is still no treatment available to patients that attenuates the observed neurological sequelae significantly. In experimental models, BDNF and anti-oxidative treatments have both shown promise, as well as inhibitors of MMPs; however, these remain unavailable for clinical testing [18]. Partly, the difficulties in finding therapeutics that reduce neurological sequelae in patients are due to the lack of understanding of the disease progression and the variability of the experimental animal models used. Our goal was, therefore, to utilize the bacteraemia-derived meningitis mouse model to describe the tempo-spatial invasion pattern of pneumococci into the brain. Furthermore, we aimed to investigate the impact of this invasion on neuronal apoptosis, microglial activation, and neurogenesis. Our results show that the pneumococci invade all brain regions in an equal manner; to our knowledge, we are the first to visualize bacterial invasion in the brain utilizing light sheet microscopy of three dimensional (3D) whole brain imaging. This bacterial invasion caused activation of microglia that showed a dynamic response depending on the symptomatology of the mice. Moreover, we observed a loss of neuroblasts in the dentate gyrus, with this niche showing an initial increase in neuroblast proliferation and migration. In summary, this study shows the dynamic changes in the brain cellular environment during pneumococcal meningitis pathogenesis over time following the progression of bacterial invasion into the brain.

## Methods

All the key resources described above in the methods are summarized in Table 1 (Key Resources Table).

### Animal experiments

Five- to six-week-old male C57BL/6 mice were supplied from Charles River and housed in a 12hr light/dark period with *ad libitum* access to water and food, in accordance with Swedish legislations (Svenska jordbruksverket, ethical permit numbers 18965-2021 and 13890-2022). We utilized a bacteremia derived meningitis model as previously described [25]. Shortly, each mouse received an intravenous injection of 1 x 10^8^ colony forming units (CFU) of *S. pneumoniae* in PBS solution through the tail vein; control animals received a PBS injection. Mice were checked for clinical symptoms every 3h and scored according to Karolinska Institutet “Assessment of health conditions of small rodents and rabbits when illness is suspected” (“Bedömning av djurhälsa för smågnagare och kanin vid misstänkt ohälsa”) template. Shortly, mice were scored 0.1 if they had one of the following symptoms: reduced activity, slow reflex, piloerection or squinty eyes. A score of 0.2 was given if the mice showed reduced activity and mobility. At each check-up, the scores were summed together. Mice were divided into three groups: 2h post infection (asymptomatic), mild symptomatic (score of 0.1/0.2) and severe symptomatic (score 0.3/0.4). Additionally, for each time point, mice were separated into three groups for downstream analysis: 1) Brain microdissection and CFU count (n=5), 2) Serial sectioning for immunofluorescence analysis (n=3) and 3) iDISCO protocol for whole brain imaging (n=1). At time of sacrifice, 5µl of blood was collected from the tail vein before the mouse was perfused with ice cold PBS through the left ventricle and the spleen collected for analysis of systemic infection; for analysis of systemic infection n=12-17. The brain was carefully collected and placed in ice-cold 4% paraformaldehyde (PFA) for downstream analysis 2 and 3. For group 1, the brain was micro-dissected into seven brain regions (striatum, hippocampus, frontal cortex, cortex, midbrain, and cerebellum). The regions were placed in ice cold PBS, weighed, and then homogenized through a 30µm cell strainer. Bacterial presence in blood, spleen, and brain was quantified by serial diluting, plating the bacteria onto blood agar plates, and CFU count.

### Preparation of bacterial strain

*Streptococcus pneumoniae* TIGR4 (serotype 4) laboratory strain and a clinical isolate serotyped 6A [9] were grown to an OD of 0.3-0.4 in Todd-Hewitt broth (THY) and stored in a solution containing 20% glycerol and THY at -80°C.

### Embryonic mouse primary neuron isolation, culture, infection, and immunofluorescence

Mouse primary cortical neurons were prepared from the cortex of E18-old embryos isolated from C57BL/6 pregnant mice as previously described [26], and in accordance to Swedish legislations (Svenska jordbruksverket, ethical permit number 17038-2020). Briefly, brains were carefully isolated from embryos, dissected and the cortical tissue triturated with fire-polished Pasteur pipettes. The isolated cell suspension was seeded on coverslips in 12-well plates (1 × 10^5^ cells/well) coated with L-ornithine. Cells were cultivated in Neurobasal™ medium (Thermo Fisher Scientific, USA) supplemented with 1% B-27 Plus Supplement (Thermo Fisher Scientific, USA) and penicillin-streptomycin 1%. Half of the media was changed every three days. Cells were cultured for 15 days, then, primary neurons were fixed with 4% PFA in PBS for 15 min at room temperature, blocked with 1% BSA in PBS for 20 min and incubated with different primary antibodies. For bacteria localization, a rabbit antiserum against serotype 4 S*. pneumoniae* capsule (SSI Diagnostica, Denmark) at 1:100 was used, followed by a goat-anti rabbit IgG Alexa Fluor 594 secondary antibody at 1:500. Slides were mounted in Prolong Gold antifade reagent (Molecular Probes, Invitrogen) and examined under a confocal microscope (Zeiss LSM900-Airy) in combination with differential interference contrast (DIC) microscopy at 63x magnification. For β-actin analysis, monoclonal IgG antibodies against β-actin (Invitrogen) and L1CAM (R&D Systems, USA) at 1:100, mouse and rabbit respectively, were used, followed by secondaries goat anti-rabbit IgG Alexa Fluor 594 and goat-anti mouse IgG Alexa Fluor 488 at 1:500. Slides were mounted in Prolong Gold antifade reagent (Molecular Probes, Invitrogen) and examined under a confocal microscope (Stellaris 5, Leica) at 63x magnification.

### Immunostaining sections

Brains were post-fixed in 4% PFA for 16 h and placed in 30% sucrose solution for minimum 4 days, before coronal sections of 30μM were cut using a microtome, and the sections frozen in a cryoprotective solution (30% sucrose, DMSO and PBS). For each analysis, three sections per mouse brain were used. The sections were washed in PBS, blocked with 2.5% BSA and permeabilized with 0.3% Triton-X-100 (Sigma) for 1hr at room temperature and stained with primary antibodies overnight at 4°C. After washing with PBS, the sections were incubated with secondary antibodies for 2hr room temperature; nuclear staining was done by adding DAPI (Abcam) for 10min. The primary antibodies used in this study were: Iba1 (Abcam, 1:1000), pneumococcal protein CCrZ (In house, 1:100), B3-tubulin (Promega,1:1000), Ki-67 (Abcam, 1:200), NeuN (,1:250) and caspase-3 (BDpharmingen, 1:250). Secondary antibodies used were Alexa Fluor conjugated anti-immunoglobulin at 1:1000 (goat anti-rabbit IgG Alexa Fluor 594 and goat-anti mouse IgG Alexa Fluor 488). The microvasculature was stained using Lectin-488 conjugate. Images was obtained taking a z-stack containing between 17-20 sections of 1µM thick. Images were taken at both 20x and 63x magnification on Zeiss LSM900-Airy, LSM800-Airy and LSM880 confocal microscope, with settings kept constant between respective image acquisition used in quantification. Images acquired for assessing caspase-3 intensity were taken on an Axio serial scanner with a magnification of 20x.

### Tissue clearing and light sheet microscopy

For tissue clearing, the iDISCO protocol was used (https://idisco.info/idisco-protocol/) [27]. The primary antibody used was pneumococcal serotype 4 anti-capsule (SSI Diagnostica, 1:250), secondary antibody used was Alexa Fluor 620 (1:1000). Whole brain images were acquired by a LaVision Biotec (UltraMicroscope II) light-sheet microscope and Imspector software. The whole brain was imaged in a total of 5mm depth, with the laser lines 488 (autofluorescence) and 639 (bacteria staining) used. The images of the autofluorescence of the brain were recorded with a resolution of x =4.797 and y = 4.796, and a z-step size of 4μm, captured with 1.26x magnification. For bacterial staining of the brain, the x and y resolution were 0.755 and a z-step size of 4μm, captured with 8x magnification. The xy images were tiled with a 10% overlap and stitched using Imspector and TeraSticher to generate 2D images.

### Image processing and analysis

Amira (ThermoFisher^TM^) was used to analyse the 3D whole brain imaging data. The autofluorescence images were sharpened and the noise reduced by using the following plug-ins: gaussian filter, unsharp masking and brightness contrast. Then, the data was segmented by applying a threshold that was determined manually. Volume rendering of the signal generated a signal of the brain, used as a “skeleton” for the subsequent overlay of bacterial signal. Images acquired from the bacterial staining were downsized to match the autofluorescence images, before the images where sharpened and background reduced by using unsharp mask and background detection correction plug in. Segmentation of the bacterial signal was done by placing a manual threshold to the images, before segmentation tool box was used to remove noise and artifacts. In all images, the outer edge of the brain was removed as this area contained a mix of high signal due to antibody unspecificities and bacterial staining (Supplemental Figure 1). The two label fields were overlaid and animated in Amira the videos were generated using DaVinci Resolve.

ImageJ (Fiji) was used to analyse the confocal and slide scanner images. To assess the intensity of Iba1 staining, the 20x magnification stacks were merged and the intensity was measured using the measure plugin of a region of interest kept constant throughout the analysis. To assess numbers of pneumococci in the cortex, we manually counted the number of CCrZ positive cells that co-localized with, or not, the microvasculature. The caspase-3 and NeuN double positive cells was counted in whole coronal sections. To assess preference of bacterial interaction with primary neurons, relative area of cell body vs. projections was measured in each field.

### Statistical analysis

GraphPad Prism-9 was used for statistical analyses (α5%). Outliers were identified by Grubb’s test. Normality was tested with D’Agostino–Pearson test. Two-way ANOVA was used to assess differences in CFU in brain areas at different groups. While Ordinary one-way ANOVA or Kruskal-Wallis test was performed with Tukey or Dunńs multiple comparisons as indicated in figure legends.

## Results

### S. pneumoniae invades the brain, equally affecting all brain regions and causing neuronal damage

The majority of pneumococcal meningitis cases follow bacterial invasion into the brain from the systemic circulation; thus, we utilized our established bacteremia-derived meningitis model to mimic the etiology of the disease [9, 13, 25]. To describe the progression of the disease, we divided mice into four groups control (PBS injected), asymptomatic, mild, and severe symptomatic. The *S. pneumoniae* strain TIGR4 was chosen to generate pneumococcal meningitis as it contains the most important virulence factors, including the pilus [28]. To evaluate the spatial invasion pattern into the brain, 3D whole brain imaging and CFU count of seven brain regions were performed. After inoculating the mice with TIGR4, the disease progressed rapidly with increased numbers of bacteria in the blood (Figure 1A) and spleen (Figure 1B) observed from the asymptomatic to severe symptomatic group. No bacteria were observed in any tissue sample taken from controls (data not shown). The number of pneumococci also increased in the brain with time, with each of the seven brain regions equally affected by the invading pneumococci as visualized by 3D whole brain imaging (Figure 1C and Supplemental Videos 1-4) and by CFU count of dissected brain regions (Figure 1D). Importantly, this invasion pattern was also observed for the piliated clinical isolate 6A; although, compared with the laboratory strain TIGR4, lower bacterial counts in the brain and reduced virulence of the strain were observed (Supplemental Figures 2A and 2B). Our results therefore indicate that piliated pneumococci invade the brain without a spatial difference, which is in unison with the bacteria employing the BBB as a primary site of invasion.

**Figure 1.**
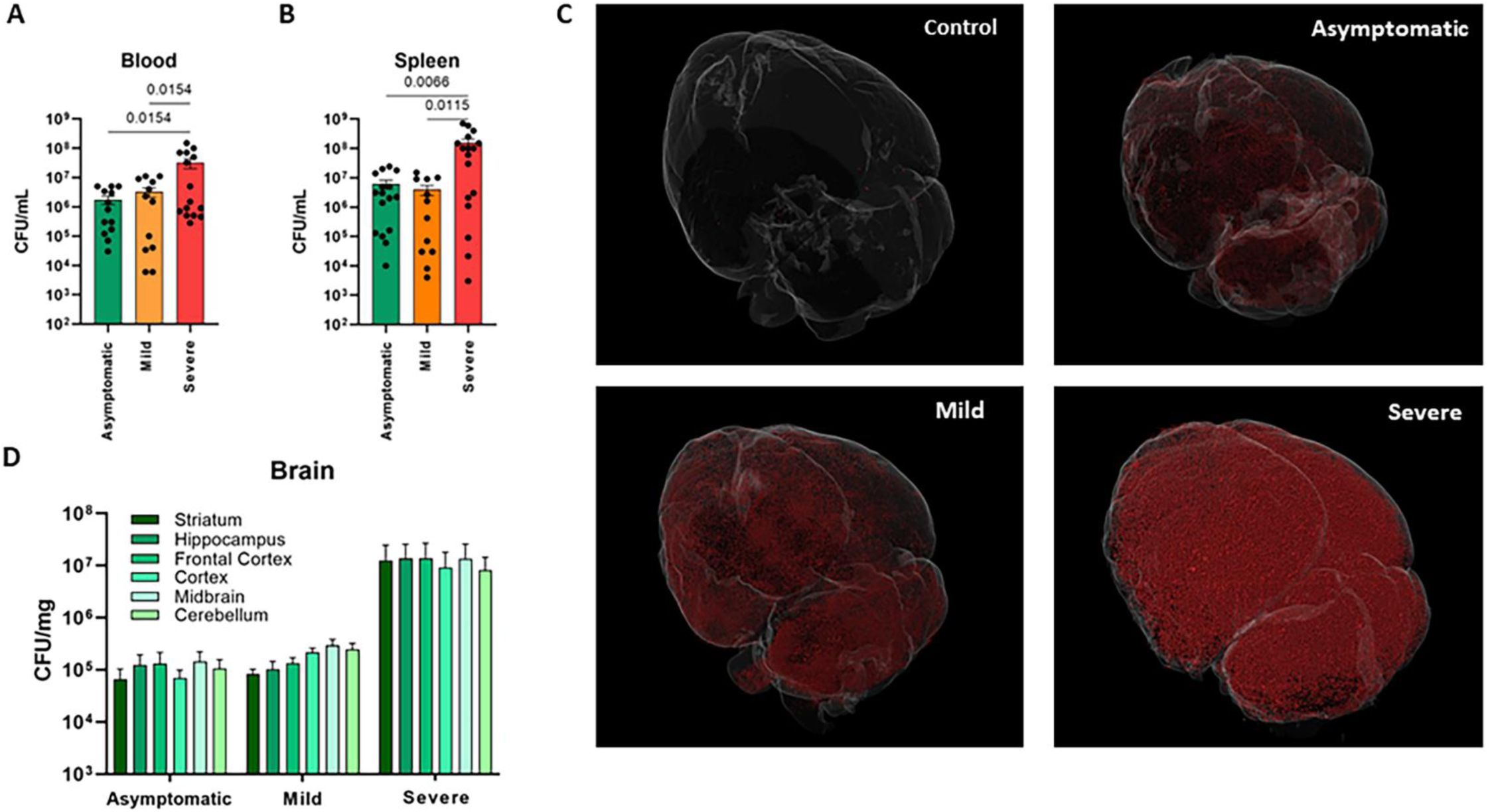
Pneumococcal invasion of the brain. **(A)** CFU in the blood increased as disease progressed after IV injection of TIGR4, n=12-15. **(B)** CFU in the spleen increased as disease progressed after IV injection of TIGR4, n= 11-17. **(C).** Pictures from 3D whole brain imaging showing a visualization of the invasion of *S. pneumoniae* (in red) into the brain, n=1. Significance was tested with one-way ANOVA and Tukey’s multiple comparison. Bars show mean ± SEM. **(D)** CFU in striatum, hippocampus, frontal cortex, cortex, midbrain, and cerebellum increased in all brain regions with time, n=5.

**Figure 2.**
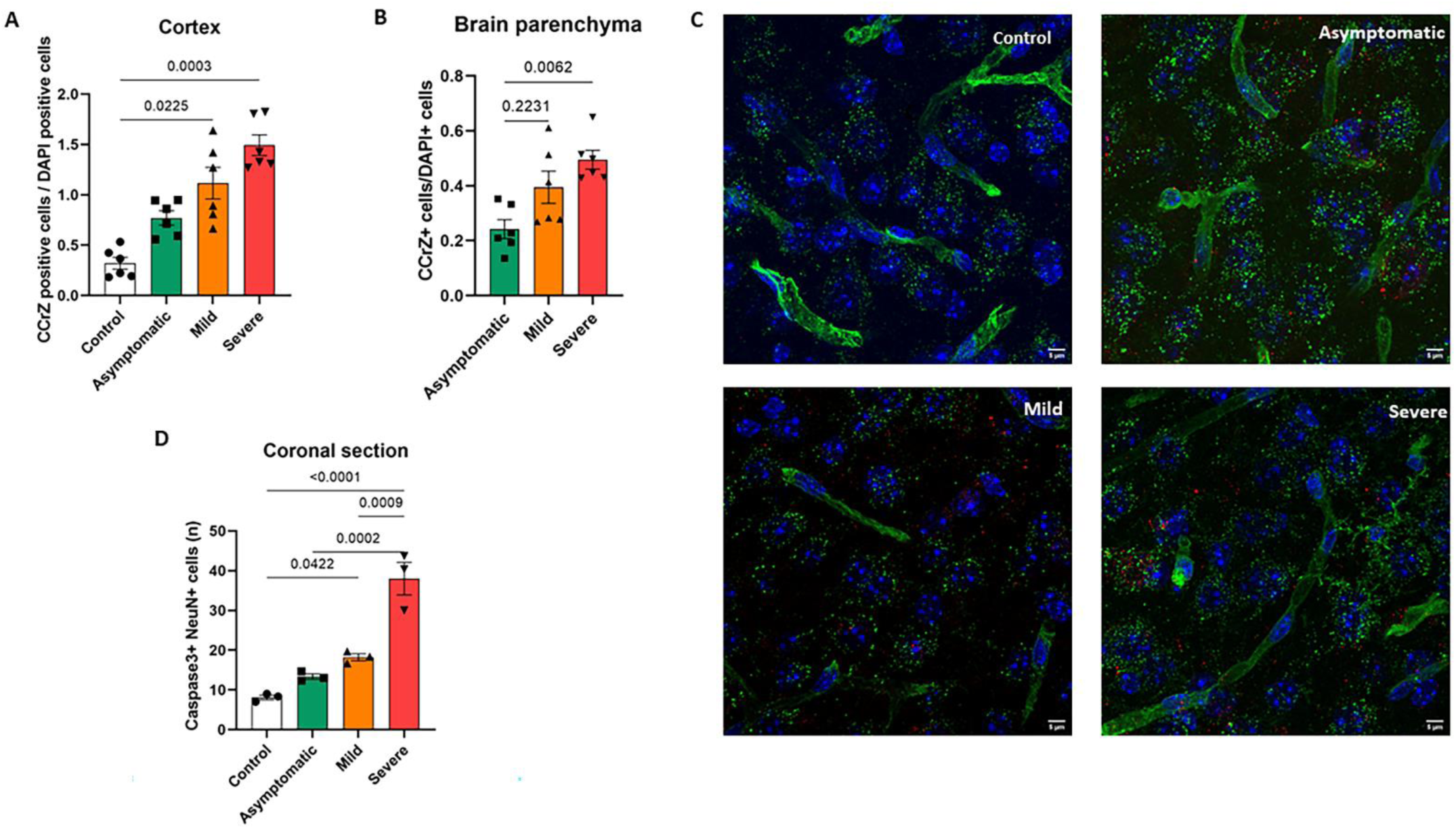
Pneumococcal invasion into the brain parenchyma causes increased caspase-3 staining. **(A)** Presence of pneumococci (CCrZ) increased in the cortex and **(B)** parenchyma during disease progression. Each dot represents the average measurement of field of view (FOV) in 3 mice, in total six FOV. **(C)** Representative images for each animal group for the quantification of pneumococci in the parenchyma and the vasculature. Bacteria stained in red, vasculature (lectin) in green and DAPI in blue. 63X magnification. Scalebar indicates 5µM. **(D)** Caspase-3 positive and NeuN positive cells increased with disease progression in the coronal sections. Each dot represent the mean value of three coronal sections for each mouse. n = 3 per animal group. Significance was tested using Kruskal-Wallis test, with Dunn’s multiple comparison or One-way ANOVA with Tukey’s multiple comparison. Bars show mean ± SEM.

Our visualization of the pneumococci using 3D whole brain imaging showed that a significant amount of bacterial signal was detected around and in the microvasculature of the brain. We therefore sought to investigate further pneumococcal invasion into the brain parenchyma. In the cortex, we observed significantly higher numbers of pneumococci with time, when comparing asymptomatic, mild, and severe mice (Figure 2A). When quantifying the co-localization between the microvasculature and pneumococci, we observed a significant increase in bacteria located in the parenchyma compared to the vasculature as disease progressed (Figures 2B and 2C). The number of vasculature-associated pneumococci also increased with time (data not shown). No difference was observed between the ratio of parenchymal/total nor vasculature/total bacteria between the three groups (data not shown), indicating that the translocation from the BBB into the parenchyma was constant and dependent on the number of total bacteria in the blood. However, in all animal groups, most bacteria were observed in the vasculature compared to the parenchyma. Bacterial infiltration into the brain caused increased caspase-3 signal in neurons, most prominent in the severe symptomatic mice, indicating caspase-3 dependent apoptosis (Figure 2D). Thus, our results indicate that the pneumococci invade into the parenchyma during all disease stages, and this is associated with a time dependent increase in apoptosis among neurons.

### S. pneumoniae preferably adheres to the cell body of primary cortex neurons

After analyzing the *S. pneumoniae* pattern of brain invasion in the different brain regions during disease’s progression, we wanted to investigate whether pneumococci have a preferential adhesion pattern on neurons at a cellular level. We have previously shown that piliated *S. pneumoniae* adheres to β-actin exposed on the cell surface of a differentiated neuron-like neuroblastoma cell line [13]. To further elaborate on this process, we investigated if TIGR4 pneumococci showed a preferential spatial binding site on primary cortex neurons. Our results indicate that TIGR4 shows a preference towards the cell body of the neuron (Figures 3A and 3B). As neuronal actin structures are notoriously different in different parts of the cell, we then assessed the presence of exposed β-actin on the plasma membrane. We saw that exposure of β-actin on the neuronal plasma membrane was enhanced after bacterial infection with a significantly increased presence in the cell body of neurons (Figures 3C and 3D), providing further indications that the pneumococci utilize β-actin, exposed on a plasma membrane damaged due to Ply and other virulence factors, to adhere and invade the neuronal cell, as we have previously speculated [10].

**Figure 3.**
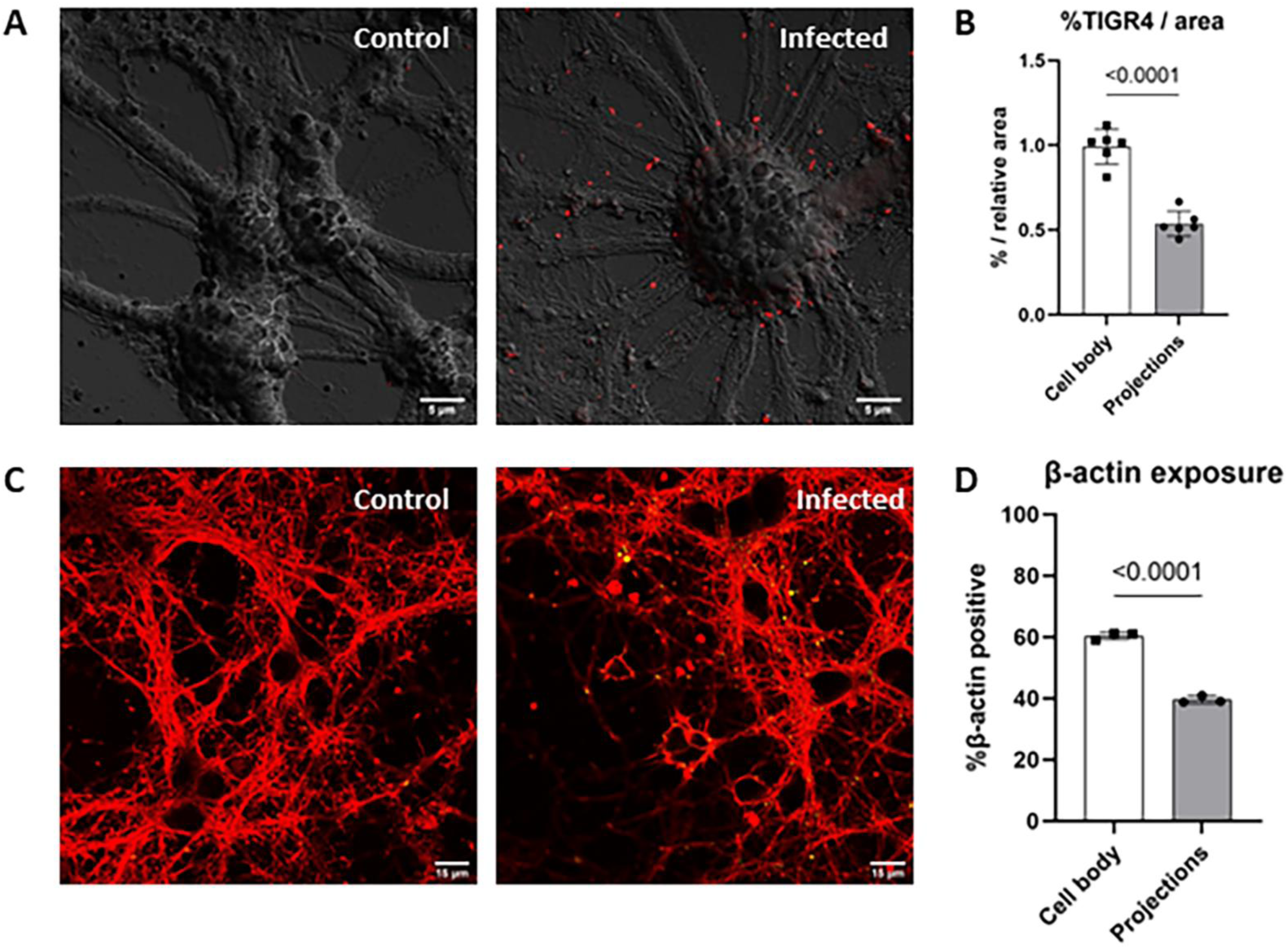
*S. pneumoniae* preferably adheres to the cell body of primary cortex neurons. **(A) (A)** Representative images of pneumococci (in red) adhered to neurons with a preference to the cell soma rather than the projections of the neurons imaged with DIC; uninfected neurons were used as control. (**B**) Quantification of bacterial presence of neuronal soma and projections; each dot represents the average measurement of one biological replicate with, at least, 5 FOV each (63x), n=6. Relative area was measured by ImageJ and used to compare results relatively. **(C)** Representative images of β-actin (in green) and its exposure on the plasma membrane stained with L1CAM (in red) after infection during TIGR4 infection and in uninfected condition. **(D)** Quantification of β-actin exposure on neuronal plasma membrane; each dot represents the average measurement of one biological replicate with, at least, 5 FOV each (63x), n=3. Data was statistical tested with an unpaired t-test. Bars show mean ± SEM.

### Pneumococcal meningitis caused a dynamic microglial response and altered the neuronal progenitor cell population in the hippocampus

Neurological sequelae in patients with pneumococcal meningitis have been linked to neuronal death in the hippocampal area [29]. Therefore, we investigated the dynamic changes of microglial activation and neurogenesis in the hippocampus during disease progression. Microglial response in the hippocampus was assessed using the Iba1 marker. The intensity of Iba1 staining decreased in asymptomatic and mild symptomatic mice in comparison to control mice (Figure 4A). Interestingly, in severe symptomatic mice, the Iba1 staining intensity was restored to levels comparable with the control group (Figure 4A). We also observed morphological differences in the four experimental groups. While the controls showed microglia with ramified processes and a small cell body, the ramifications were reduced in the three other groups, with a morphology showing more dense protrusions and a more amoebic shape (Figure 4B). Thus, our results indicate that the decrease in Iba1 staining intensity observed in asymptomatic and mild symptomatic mice could be due retraction of microglia protrusions. Based on our findings, we can hypothesize that the increase in Iba1 staining observed in the severe symptomatic mice is due to infiltration of peripheral macrophages, which also express the Iba1 marker.

**Figure 4.**
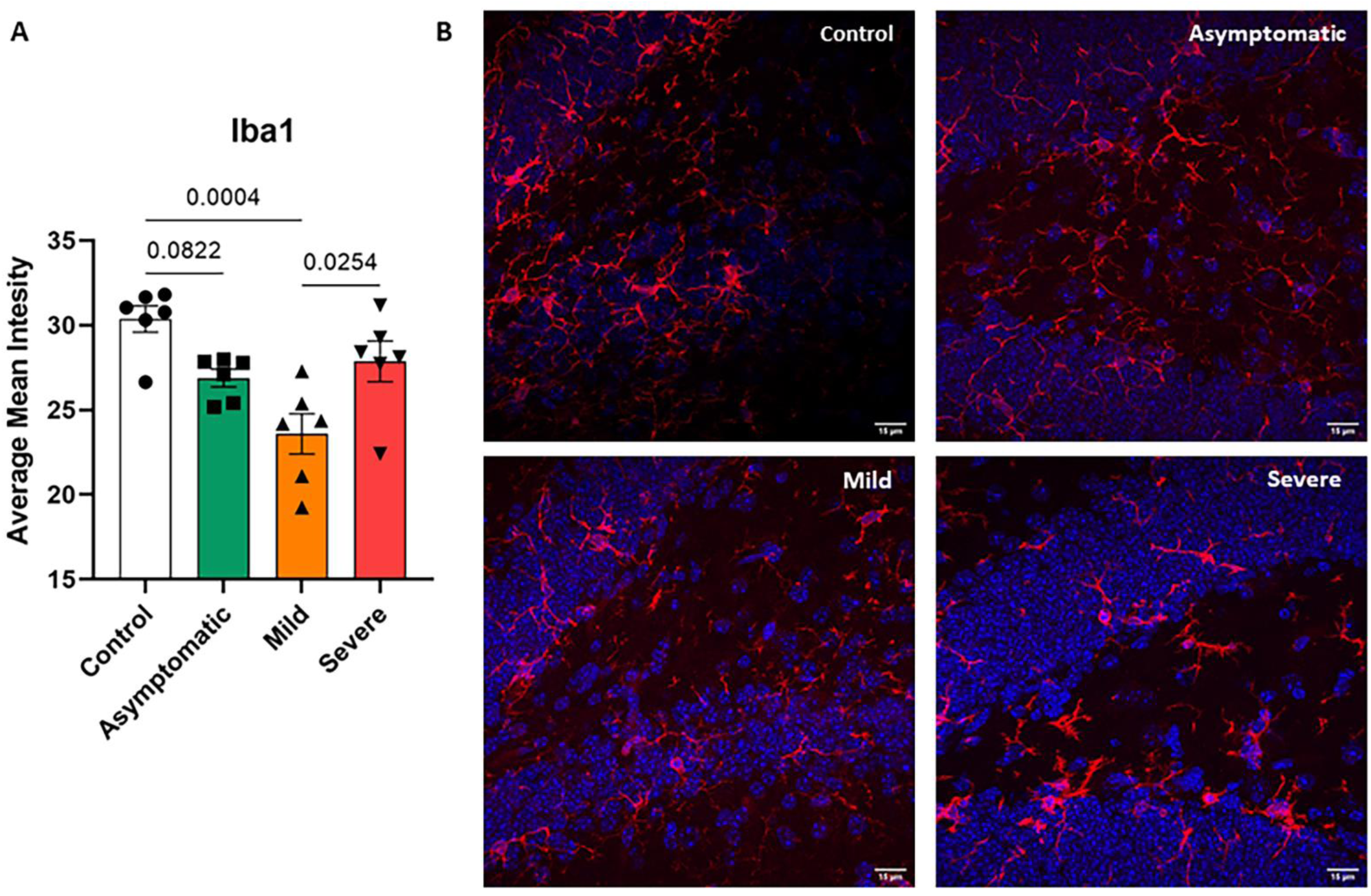
Microglial activation in the hippocampus during disease progression. **(A)** The average mean intensity of Iba1 decreased in the early stages of disease progression, then increased in the later stage. Each dot represents the average measurement of FOV in 3 mice, in total six FOV. n=3. **(B)** Representative images from each mouse group. As disease progress the microglia show an increase in cell body and decreased ramifications, typical of an activated phenotype. Microglia as shown in red and DAPI in blue. White arrows indicate activated phenotype. Magnification 63X, Scalebar indicates 15µM. Significance was tested using a one-way ANOVA, with Tukey’s test for multiple comparison. Bars show mean ± SEM.

Alteration in microglia activation has been shown to influence the neurogenesis in the dentate gyrus of the hippocampus [30]. We used the neuroblast marker B3-tubulin and proliferation marker Ki-67 to evaluate how neurogenesis was affected during disease progression. We observed a decrease in B3-tubulin staining in the SGZ in the dentate gyrus. In contrast, this staining was increased in the hilus (Figures 5A-C), suggesting that neuroblasts are decreasing in numbers and/or migrating towards neuronal circuits. Ki-67 is upregulated in newly proliferated cells and thus, combined with B3-tubulin positive marker, shows evidence of neuroblast proliferation. When quantifying the number of double-positive cells, we observed an initial increase in numbers from control animals to asymptomatic and mild symptomatic mice; however, in severe symptomatic mice, the numbers of double-positive cells decreased and were comparable with control animals (Figures 5C and 5D). This would indicate that the niche was trying to compensate for the loss of neuroblasts and/or loss of neurons in the area and tried to regenerate, before failing to do so due to the continuous pro-inflammatory and cytotoxic environment created by pneumococcal products.

**Figure 5.**
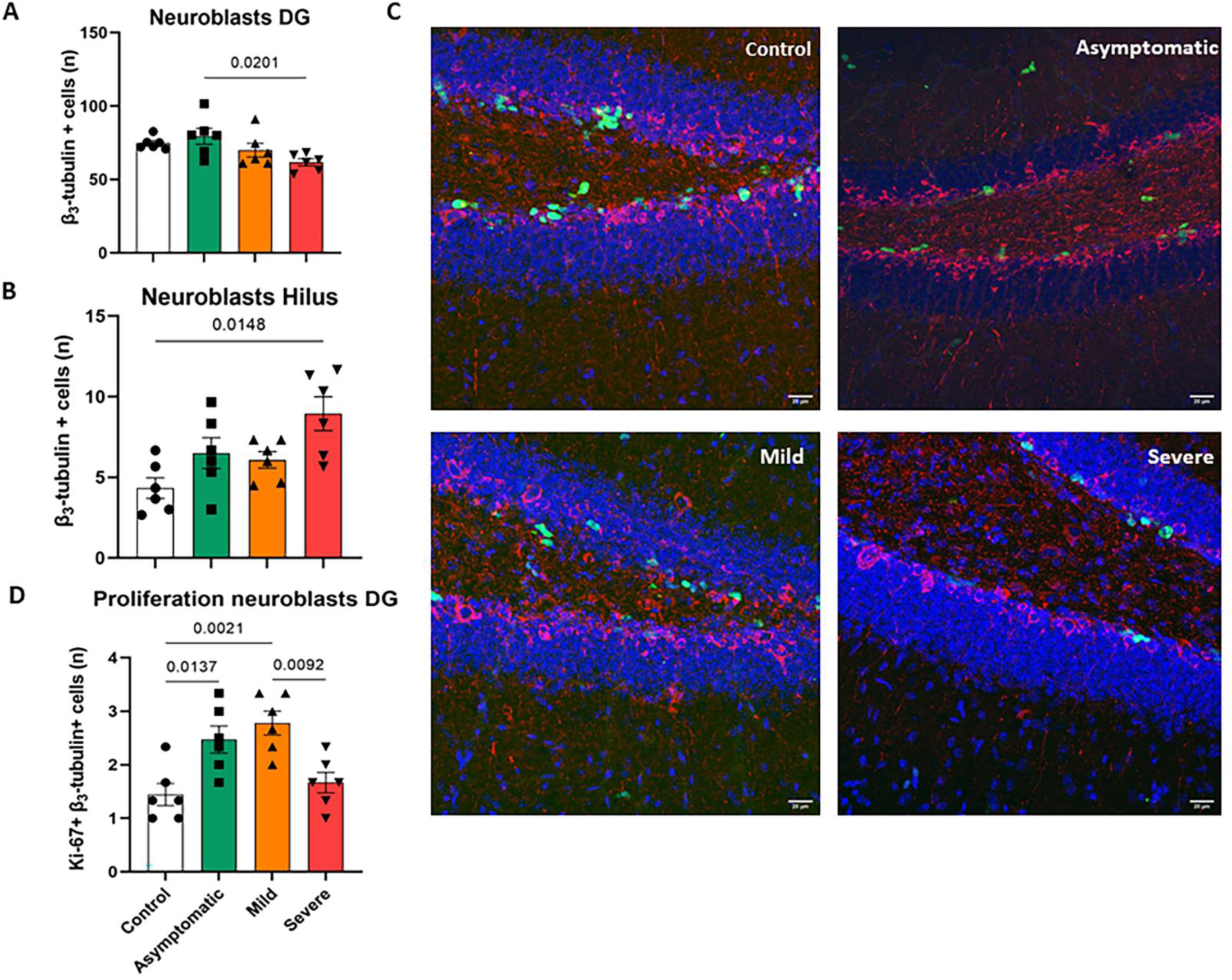
The dynamic cellular response of the neurogenic niche in the hippocampus. **(A)** Quantification of β3-tubulin positive cells in the dentate gyrus. **(B)** Quantification of β3-tubulin positive cells in the hilus. **(C)** Quantification of β3-tubulin and Ki-67 double positive cells in the dentate gyrus. **(D)** Representative images from each mouse group, DAPI in blue, Ki-67 in green and β3-tubulin in red. Magnification 63X, Scalebar indicates 20µM. Significance was tested using Kruskal-Wallis test, with Dunn’s multiple comparison. Bars show mean ± SEM.

## Discussion

The World Health Organization (WHO) defines meningitis as “devastating” because, even if the bacterial infection is adequately cured, permanent neurological disabilities occur in approximately half of survivors https://www.who.int/news-room/fact-sheets/detail/meningitis. These long-term neurological sequelae are a burden for patients themselves, but also for health care systems worldwide [31]. Importantly, in retrospective studies, children diagnosed and surviving bacterial meningitis show reduced memory and learning abilities in school and throughout life [32–34]. This highlights the importance of a better understanding of the pathophysiological process during bacterial meningitis. In this study, focused on pneumococcal meningitis, we have described this pathological process in a dynamical way, and we have shown that deleterious events occur early in disease progression with all brain regions affected by pneumococcal invasion.

To our knowledge, we are the first to combine the use of cleared brain tissue and light-sheet microscopy to visualize bacteria in the brain in 3D whole brain imaging; this, together with counting of CFU numbers from brain microdissection, showed that pneumococci invade uniformly into the parenchyma. This heterogenous brain invasion reflects the clinical reality of the neurological sequelae post bacterial meningitis, as the vast variety of sequelae observed are due to damaged motor, sensorial or emotional functions which are controlled by different brain regions [29, 30]. Furthermore, in line with previous research from rodent and zebrafish models, the pneumococci were shown to adhere and to be in close proximity with blood vessels, indicating this as the primary site of invasion [11, 35]. We observed a high number of *S. pneumoniae* in the brain shortly after inoculation of the mice, with bacteria already invading into the parenchyma two hours after infection. While bacterial invasion of the brain progressed, neuronal death also increased accordingly; neuronal apoptosis has been shown to be partially caspase-3-dependent, and an increase in caspase-3 was also observed in our study [20].

Our results support the concept that, once present in the systemic circulation, the bacteria do have the capacity to cross the BBB immediately, and that symptoms of disease are not a consequence of bacterial numbers in the brain, but rather of the subsequent neuroinflammatory response. As expected, and in line with the literature, the pneumococcal invasion caused a morphological change in the microglial cell population, which showed retraction of protrusions and a denser cell body indicating an activated state [36–38]. Iba1 is widely used to stain microglia, although its exact function is not completely known and, in brain diseases, it has been reported to be both increased and decreased in intensity [39]. Interestingly, we show that the Iba1 intensity showed a temporal dynamic change, with lower levels observed for the asymptomatic and mild symptomatic group, compared to the control group. However, the severe group showed markedly increased Iba1 staining compared to all the other three animal groups. We therefore hypothesize that this shift in the staining intensity is due to the morphological changes in the microglia, and a potential influx of peripheral macrophages into the brain. Inflammation and microglial activation are detrimental for the generation of new neurons in the dentate gyrus of the hippocampus [30, 40]. In post-mortem brains from bacterial meningitis cases, an increase in neural progenitors was reported [41]. These progenitors can differentiate into either astroglia or neuroblasts; and in rodent models of meningitis, a decrease in neuroblasts has been reported [42, 43]. Our results are in line with these previous findings, in fact, here we show an overall loss of neuroblasts in the dentate gyrus. Additionally, it seems that the neurogenic niche tries to compensate for a loss of neurons in the early stage of disease, causing an increased proliferation of neuroblasts which is inhibited as the disease progresses. This loss has been shown to withstand for weeks, causing a reduction in neurons, and potentially causing cognitive deficiencies [44, 45].

The actin cytoskeleton has been linked to bacterial pathogenesis in several ways, from invasion, intracellular motility and navigation to phagocytosis escape [46, 47]. Ply has been previously shown to cause dendritic spine collapse via de formation of membrane pores that disrupt the critical cytoskeletal structures that sustain such spines, many of them depending on F-actin processes [48]. We have previously suggested a synergy between RrgA and Ply in pneumococcal interaction with neurons; in fact, using purified proteins we have shown that, once in contact with neuron-like cells, Ply alters the plasma membrane increasing the exposure of β-actin, which can therefore be bound by pneumococci, therefore promoting bacterial adhesion [10]. Here, we show that the infection with *S. pneumoniae* causes an increase of β-actin exposure on the plasma membrane of primary murine cortical neurons. Moreover, confocal microscopy analysis revealed that pneumococcal adhesion preferentially occurred on the cell body of neurons rather than their projections, axon- and dendrite-like structures, most likely because the exposure of β-actin on neuronal plasma membrane was significantly higher neuronal soma rather on the projections, providing further confirmation on the mechanism of pneumococcal invasion to neurons.

While several studies have investigated and described the deleterious events in the brain cellular environment during bacterial meningitis pathogenesis, the majority utilizes an intracisternal model [49–51]. This model has several benefits, including the possibility to use strains without good capability to penetrate the BBB and is, furthermore, the only way to control the exact number of bacteria that enters the central nervous system. However, precisely because it bypasses the initial crossing of the BBB, it does not properly mimic the effect of a systemic infection on the pathological process. In contrast, the bacteraemia-derived meningitis model utilized in this study considers how the progression of the disease occurs in most clinical cases. Furthermore, the effect of systemic infections on the brain environment, and the fact that pneumococci cause damage to the hippocampus before getting access into the brain, underlines the strength of using this model when evaluating the invasion and dynamic cellular changes in the brain environment. As mentioned, here we provide here a clear visualization of the invasion pattern of pneumococci into the brain using light-sheet microscopy. Light-sheet microscopy combined with 3D brain imaging has been applied before in literature, however always investigating at specific cells in certain brain regions, such as hippocampal neurons in mouse models [52, 53]. A step forward in terms of spatial completeness was done by Ahrens et al that investigated communication between large population of neurons in a larval zebrafish model via a more comprehensive whole brain imaging [54]. More recently, studies by Cong et al and Zhang et al have adopted whole brain imaging to investigate different neural structures and their respective activities [55, 56]. Our study shows the use of light-sheet microscopy combined with whole brain imaging to monitor invasion of the brain by blood-borne bacterial pathogens during the progression of an invasive infectious disease. Although the bacterium has an average size of 1µm, we could clearly see bacterial signal in the brain tissue. However, we find it unlikely that these pneumococci represent single bacterium, as this would require higher magnification, currently not available for light-sheet microscopy. Despite this, we believe that this technique will be useful for future investigations of the brain invasion mechanism of invasive pneumococci and other pathogens.

In conclusion, by combining a bacteremia-derived meningitis mouse model and light-sheet microscopy for 3D whole brain imaging, we have provided a comprehensive spatio-temporal analysis of brain invasion by blood-borne *S. pneumoniae*, a crucial process in the pathogenesis of pneumococcal meningitis. In addition, this combination of *in vivo* model with high-resolution microscopy analysis allowed us to study the inflammatory, neuronal apoptotic and neurogenic response, from the time when pneumococci start entering the brain until the acute phases of disease in which the brain is severely invaded by the pathogen.

## Supporting information

Key Resources Table

## Acknowledgments

The authors sincerely thank Dr. Shigeaki Kanatani from the Biomedicum Imaging Core (BIC) Facility at Karolinska Institutet for the assistance during the light-sheet microscopy and 3D whole brain imaging and downstream analysis, Prof. Jan Willem Veening (Department of Fundamental Microbiology, University of Lausanne, Switzerland) for kindly providing the anti-CcrZ antibody, and Prof. Carlos Orihuela (Department of Microbiology, University of Birmingham, Alabama USA) for kindly providing the *S. pneumoniae* TIGR4 strain. This research was supported by The Swedish Research Council (Vetenskapsrådet), Bjarne Ahlström Memorial Fund, The European Society of Clinical Microbiology and Infectious Diseases (ESCMID), Wera Ekström Fund for Pediatric Research, Magnus Bergvall Foundation, and Tore Nilson Foundation.

## Author contributions

FI and KF designed the study. KF and MTV performed the experiments, analysed data. KF, MTV and FI wrote the manuscript. KW and LO participated in the experimental approach and revised the manuscript.

