## Supplementary Information for "Spatio-temporal brain invasion pattern of *Streptococcus pneumoniae* and dynamic changes of the cellular environment in meningitis pathogenesis"

### Supplemental Information

**Supplemental Figure 1. Detection of bacterial signal in Amira. (A)** Representative images showing the bacterial infiltration into the brain in each animal group, and the subsequent signal detected by applying a threshold. Magnification 12.6X. **(B)** The bacterial signal (in blue) was detected by a threshold set to each brain; representative pictures are shown for each animal group.

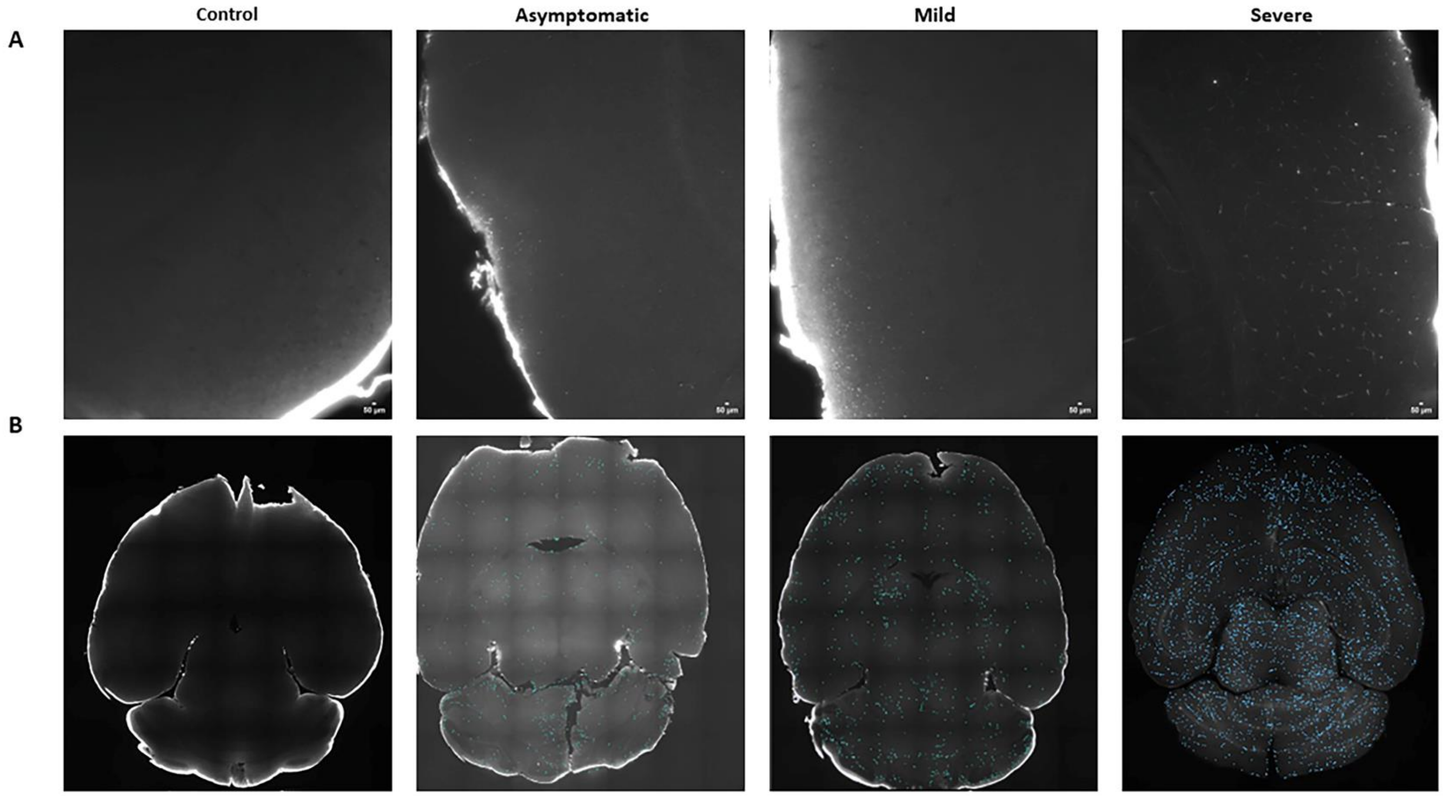

**Supplemental Figure 2. Pneumococcal invasion in the brain of meningitis clinical isolate serotype 6A. (A)** Increase in CFU in the blood as disease progresses after IV injection of TIGR4. **(B)** The CFU in striatum, hippocampus, frontal cortex, cortex, midbrain, and cerebellum increased in all brain regions with time.

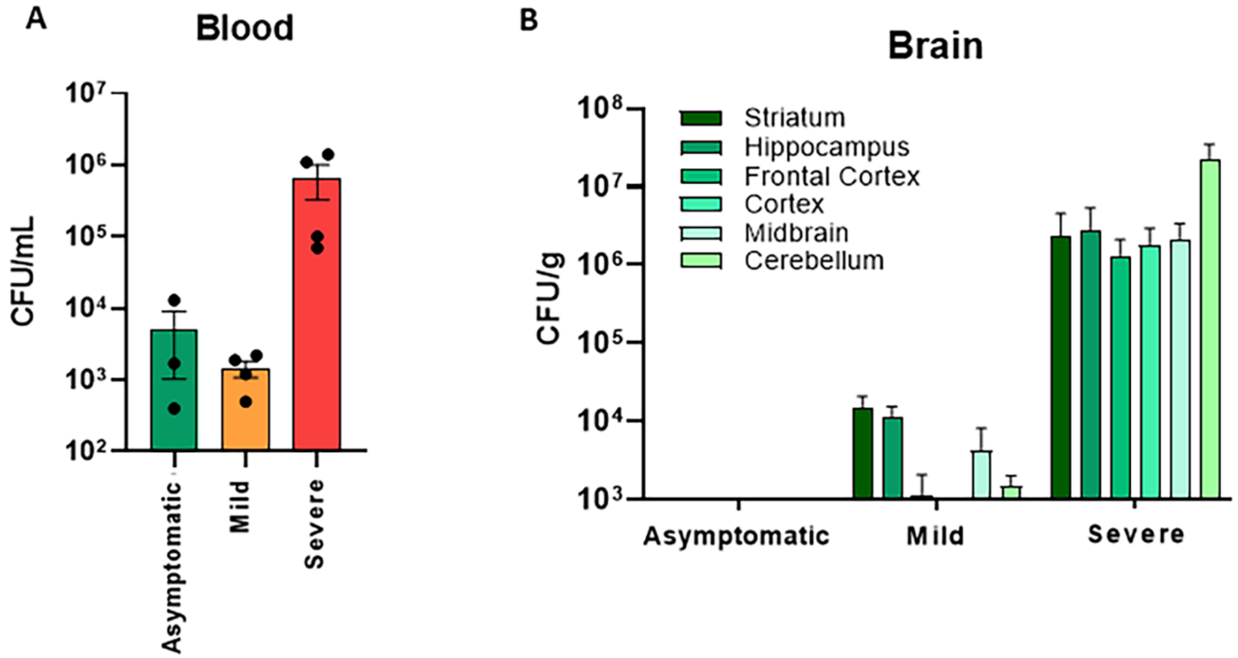


**Supplemental Videos. Invasion of the brain by *S. pneumoniae* TIGR4 visualized via 3D whole brain imaging.** Invasion of pneumococci (in red) invading the brain visualized by whole brain imaging in the asymptomatic **(1)**, mild **(2)**, severe **(3)** experimental groups; the brain from an uninfected mouse was also imaged as control **(4)**; n=1 per each group.
