## Supplementary material for "Spatio-temporal brain invasion pattern of *Streptococcus pneumoniae* and dynamic changes of the cellular environment in meningitis pathogenesis": Key Resources Table

**Table 1. Key resources table**

| REAGENT or RESOURCE | SOURCE | IDENTIFIER |
|---|---|---|
| Antibodies | | |
| Pneumococcus Type 4 serum, rabbit polyclonal | SSI Diagnostica | Catalog: 16747, Lot: G416C1 |
| Anti-Iba1, goat polyclonal | Abcam | Catalog: ab5076, Lot: 1002201-2 |
| Anti-βIII Tubulin, mouse monoclonal | Promega | Catalog: G7121 |
| Anti-Ki67, rabbit polyclonal | Abcam | Catalog: ab15580 |
| Anti-NeuN, mouse monoclonal | Merck Millipore | Catalog: MAB377, Clone: A60 |
| Caspase 3, rabbit monoclonal | BD Pharmigen | Catalog: 700182, Clone: 9H19L2 |
| Anti-rabbit Alexa Fluor 594, goat polyclonal | Invitrogen | Catalog: 10266352 |
| Anti-mouse Alexa Fluor 488, goat polyclonal | Invitrogen | Catalog: A28175 |
| Anti-β-actin, mouse monoclonal | Invitrogen | Catalog: AM4302, Clone: AC-15 |
| Anti-L1CAM, rabbit monoclonal | Abcam | Catalog: ab208155, Clone: EPR18750 |
| Anti-rabbit Alexa Fluor 647, goat polyclonal | Invitrogen | Catalog: A21244, Lot: 2326489 |
| Anti-CcrZ, rabbit polyclonal | Kind donation by Jan Willem Veening's group (Department of Applied Microbiology, University of Lausanne, Switzerland) | |
| Donkey Serum | Jackson Immunoresearch | 017-000-121 |
| Bacterial and Virus Strains | | |
| *Streptococcus pneumoniae:* TIGR4: WT | Kind donation by Carlos Orihuela's group (Department of Microbiology, University of Birmingham, Alabama USA) | Taxonomy ID: 1313 |
| *Streptococcus pneumoniae*: Serotype 6A meningitis clinical isolate | Pneumococcal strain collection of Federico Iovino's group, originally a kind donation from Diederik van de Beek's group (Department of | Taxonomy ID: 1313 |

| | | |
|---|---|---|
| | Neurology, Amsterdam Academic Medical Center, Netherlands) | |
| Chemicals, Peptides, and Recombinant Proteins | | |
| Isoflurane | Baxter | Catalog: HDG9623V |
| Triton X100 | Sigma | Catalog: X100-500ML |
| Tween-20 | Sigma | Catalog: P9416-100ML |
| DMSO | Fisher | Catalog: D128-4 |
| Sodium Azide | Sigma | Catalog: 58032-100G |
| Glycine | Sigma | Catalog: G7126-500G |
| Heparin | Sigma | Catalog: H3393-50KU |
| Methanol | Fisher | Catalog: A412SK-4 |
| Hydrogen Peroxide 30% | Sigma | Catalog: 216763-100ML |
| DiChloroMethane | Sigma | Catalog: 270997-12X100ML |
| DiBenzylEther | Sigma | Catalog: 108014-1KG |
| Paraformaldehyde 4% | HistoLab | Catalog: HL96753.1000 |
| Bovine Serum Albumin | Sigma | Catalog: A3294 |
| Experimental Models: Organisms/Strains | | |
| Mus musculus: strain C57BL/6: wt | Charles River | Taxonomy ID: 10090 |
| Software and Algorithms | | |
| Fiji ImageJ | National Institutes of Health, USA | Version 2 |
| Prism | GraphPad | Version 9 |
| Amira 3D | ThermoFisher | Version 2023.1.1 |
| DaVinci Resolve | Black Magic Design | Version 18 |
| Imspector | Abberior Instruments | Version 7 |
| TeraStitcher | Github, free software developed by A. Bria and G. Iannello | v1.10.12 |
